# E-cigarette aerosol exacerbates cardiovascular oxidative stress in mice with an inactive aldehyde dehydrogenase 2 enzyme

**DOI:** 10.1101/2021.11.02.466292

**Authors:** Xuan Yu, Xiaocong Zeng, Feng Xiao, Ri Chen, Pritam Sinharoy, Eric R. Gross

**Author notes:** **Address Correspondence to**: Eric R. Gross, MD, PhD, Department of Anesthesiology, Perioperative and Pain Medicine, School of Medicine, Stanford University, Stanford, CA. 94305.

## Abstract

**Aims:** E-cigarette aerosol containing aldehydes, including acetaldehyde, are metabolized by the enzyme aldehyde dehydrogenase 2 (ALDH2). However, little is known how aldehyde exposure from e-cigarettes, when coupled with an inactivating ALDH2 genetic variant, ALDH2*2 (present in 8% of the world population), affects cardiovascular oxidative stress. The aim of this study was to determine how e-cigarette aerosol exposure, when coupled with genetics, impacts cardiovascular oxidative stress in wild type ALDH2 and ALDH2*2 knock-in mice.

**Methods and Results:** Using selective ion flow mass spectrometry, we determined that e-cigarette aerosol contains acetaldehyde that are 10-fold higher than formaldehyde or acrolein. Next, using wild type ALDH2 and ALDH2*2 knock-in rodents, we identified organ-specific differences in ALDH2 with the heart having ~1.5-fold less ALDH2 enzyme activity relative to the liver and lung. In isolated cardiac myocytes, acetaldehyde exposure (30 seconds, 0.1-1 μM) caused a 4-fold greater peak in calcium levels for ALDH2*2 relative to ALDH2 cardiomyocytes. ALDH2*2 cardiomyocytes exposed to acetaldehyde also demonstrated a 2-fold increase in ROS production and 2.5-fold increase in 4HNE protein adducts relative to ALDH2 cardiomyocytes. For intact rodents, ALDH2*2 knock-in mice exposed to e-cigarette aerosol had an increased heart rate beginning 5 days after exposure compared to wild type ALDH2 mice (775±30 bpm versus 679±33 bpm, respectively, \**p*<0.01, n=7-8/group). E-cigarette aerosol exposure also exacerbated oxidative stress in ALDH2*2 heart homogenates, including a 1.3-fold higher protein carbonyl level, a 1.7-fold higher lipid peroxide level and 1.5-fold greater phosphorylation of NF-κB relative to wild type ALDH2 homogenates.

**Conclusions:** The increased oxidative stress from e-cigarette aerosol aldehydes triggers the proinflammatory NF-κB pathway. As ALDH2 expression and activity is lower in the heart relative to the lung, the heart could be more susceptible to increases in cardiovascular oxidative stress from e-cigarette aerosol; particularly for those carrying an ALDH2*2 genetic variant which limits acetaldehyde metabolism.

**Graphical Abstract:** 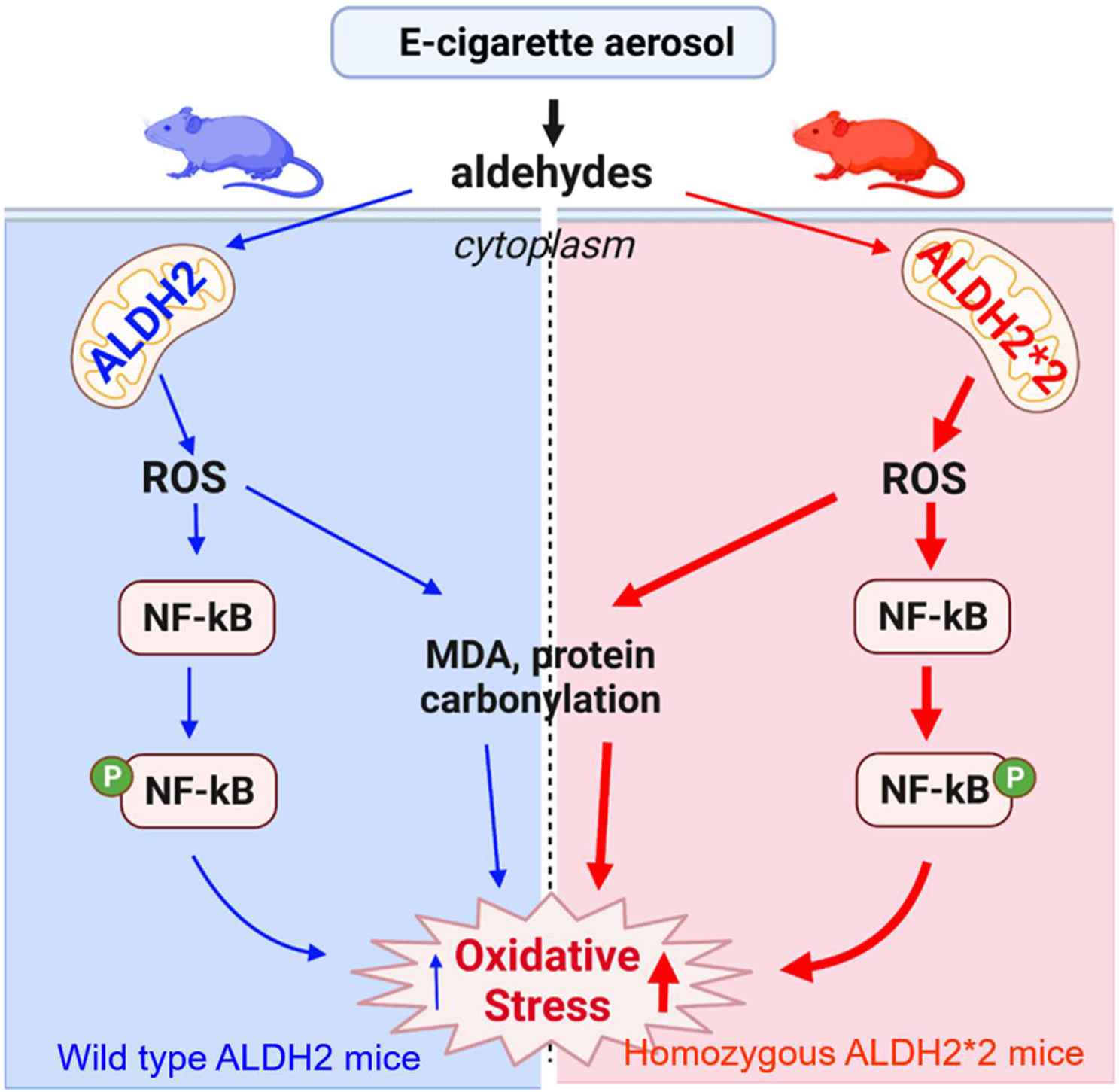

E-cigarette aerosol exposure triggers increases in ROS that lead to increased protein carbonylation, MDA production, and elevates phosphorylated NF-kB. This exposure to e-cigarette aerosol leads to increases in cardiovascular oxidative stress. For the ALDH2*2 variant, there is limited ability to metabolize the reactive aldehydes from e-cigarette aerosol and with increased levels of oxidative stress at baseline relative to wild type ALDH2, e-cigarette aerosol increased oxidative stress, protein carbonylation, and phosphorylation of NF-kB relative to wild type ALDH2 rodents.

## 1. Introduction

Electronic cigarettes (e-cigarettes) are increasing in popularity specifically among teens and young adults [1]. E-cigarette aerosol, although containing less chemicals compared to conventional tobacco cigarette smoke, contains aldehydes produced from the combustion of propylene glycol (PG) and vegetable glycerin (VG) within the e-liquid [2]. Importantly, aldehyde exposure is estimated to contribute to over 92% of the cardiopulmonary disease risk from tobacco smoke [3] and recognized by the Institute of Medicine as one of the most significant cardiovascular toxins within tobacco smoke [4]. However, whether the aldehydes within e-cigarette aerosol are detrimental to the cardiovascular system has not been extensively studied.

Generally, aldehydes are deleterious to the cell and can form aldehyde-induced protein adducts leading to cellular dysfunction and cell death [5]. These aldehydes are metabolized by the enzyme, aldehyde dehydrogenase 2 (ALDH2). In particular, ALDH2 has a Km for acetaldehyde that is 900-fold lower relative to cytosolic ALDH1 making ALDH2 selective and specific for acetaldehyde metabolism [6]. In the cardiovascular system, ALDH2 activation is important in limiting mast cell renin release and mitigating cellular injury from ischemic events by protecting the heart from aldehyde-induced injury [7, 8].

However, an inactivating ALDH2 genetic variant, known as ALDH2*2, severely limits aldehyde metabolism. The ALDH2*2 genetic variant is present in approximately 30% of people of East Asian descent (540 million people or 8% of the world population) [9]. This genetic variant leads to an increased cardiomyocyte cell death during ischemia-reperfusion injury [10–13], increased risk of coronary artery disease [14, 15], as well as alcohol-induced heart disease [16]. However, it is unknown how aldehydes present within e-cigarette aerosol impact the cardiovascular system. This is particularly important since Asian-American users of e-cigarettes, particularly among young adults [17, 18], have increased in recent years, from 2% reported in 2013 to 10% in 2018 [19]. For these reasons, the aim for this study is to determine whether in ALDH2*2 variant mice e-cigarette exposure exacerbates cardiovascular oxidative stress relative to wild type ALDH2 mice.

## 2. Methods

### 2.1. Quantification of aldehydes and nicotine in e-cigarette aerosol

All products tested were purchased online or from local vape shops including Blu, Halo and JUUL e-cigarettes. The chemical composition for each electronic cigarette tested is listed (Table 1). Aerosols were generated from electronic cigarettes by drawing from an e-cigarette connected to a silicone tubing using a Masterflex peristaltic pump (model 7518-00, Cole-Parmer Instrument company, Chicago, IL) set at a rate of 10 mL/sec. Approximately 0.4 L of e-cigarette aerosol was collected into 0.5 L Tedlar Bags (Zefon International, Ocala, Florida). For each e-cigarette, aerosol collection and measurements were performed 20 times. Aldehydes and nicotine levels in e-cigarette aerosol were quantified immediately after collection in real time using selective ion flow tube mass spectrometry (SIFT-MS, Syft, Christchurch, New Zealand). The reagent ions, reaction ratio, and mass used to detect nicotine and aldehydes by SIFT-MS are summarized (Supplemental Table 1).

**Table 1.** E-cigarette chemical composition. Three e-cigarette brands were quantified for aerosolized nicotine and aldehyde content including Blu, Halo and Juul. Brand name, chemical composition on the packaging insert, and nicotine concentration are listed.

| **Brand Name** | **Listed Ingredients** | **Flavor** | **Nicotine (mg/mL)** |
| --- | --- | --- | --- |
| **Blu Plus+ Rechargeable** | Vegetable Glycerin, Propylene Glycol, Nicotine, Natural and Artificial Flavors, Water | Classic Tobacco | 24 |
| **Halo G6 Rechargeable** | Propylene Glycol, Glycerin, Flavorings, Nicotine | Prime 15 | 24 |
| **Juul** | Glycerol, Propylene Glycol, Flavor, Nicotine, Benzoic acid | Classic Tobacco | 50 |

### 2.2. ALDH2*2 knock-in mice

In this study, we initially measured ALDH2 expression for both male and female rodents. Based on these findings and since men tend to use tobacco products more than women (18% of men smoke e-cigarettes compared to 12% of women [20]), we used male mice for the *in vivo* e-cigarette exposure and *in vitro* cell based studies. In addition, using only male mice reduces the variability and increases the power of this study to comply with the 3Rs for animal use.

To study the effects of e-cigarettes on the ALDH2*2 variant, we used an ALDH2*2 knock-in mouse on a C57/BL6 background generated using homologous recombination [21]. We created the mouse by knocking out the wild type ALDH2 gene followed by editing and inserting a single copy of the ALDH2*2 variant gene (E504K). The ALDH2*2 mice closely resemble characteristics of humans carrying the ALDH2*2 genetic variant. Particularly when challenged with alcohol, the mice accumulate acetaldehyde similar to the levels seen in humans carrying the ALDH2*2 genetic variant [21].

For this study, male and female 8-10 week old mice were initially used to characterize ALDH2 expression for heart, lung and liver. Further studies were then performed using 8-10 week old male homozygous ALDH2*2 mice and age-matched male wild type ALDH2 mice. Mouse exposure to e-cigarette aerosol or room air were carried out by ZX and XY, and data analysis occurred blinded by XY and FX.

All animals were maintained in a constant 12-h dark/12-h light cycle in an AAALAC-accredited Veterinary Service Center at Stanford University. Food and water were available ad libitum. All animal procedures were performed in accordance with National Institutes of Health guidelines for the humane care of laboratory animals and approved by the Stanford University Administrative Panel on Laboratory Animal Care (APLAC). In accordance with approved guidelines, all mice were euthanized by intraperitoneal injection of 100 mg/kg inactin hydrate (Sigma), and euthanasia were confirmed with failure of responding to toe pinch.

### 2.3. ALDH2 expression and ALDH activity assay in organ homogenates

Organs including heart, lung and liver from unexposed wild type ALDH2 mice and ALDH2*2 mice were harvested, homogenized and centrifuged at 10000 rpm for 10 minutes at 4°C (in mannitol-sucrose buffer: 210 mM mannitol, 70 mM sucrose, 5 mM MOPS, and 1 mM EDTA, pH 7.4). Protein concentration was determined by Pierce BCA protein assay kit (Thermo Scientific). The ALDH2 protein expression for heart, lung, and liver homogenates were analyzed by western blot as described previously [22]. Samples were resolved on 10% SDS-polyacrylamide gels, transferred onto PVDF membrane and incubated with ALDH2 primary antibody (Abcam 1:1000) overnight. This was followed by incubation with anti-goat secondary antibody (Invitrogen 1:2000), and were incubated for chemiluminescence at room temperature for 5 minutes and imaged by using an Azure cSeries gel imaging system (Azure Biosystems, Dublin, CA).

ALDH enzymatic activity was measured as described [8]. Briefly, 200 μg of mitochondrial fraction was incubated with 2.5 mM cofactor NAD+ and 10 mM acetaldehyde, and the increase in NADH production was measured over time by spectrophotometer (DU800, Beckman Coulter, Indianapolis, IN) at 340 nm wavelength. The ALDH2 enzymatic activity was presented as μmol NADH /min/mg protein.

### 2.4. Primary mouse cardiac myocyte studies

Adult male cardiac myocytes were isolated from wild type ALDH2 and ALDH2*2 mice as previously described [23]. Briefly, hearts were rapidly excised and washed with calcium free Krebs-Henseleit perfusion buffer. After cannulating the aorta, blood was flushed and the hearts were subjected to retrograde perfusion at 37°C with perfusion buffer. After 4 minutes of perfusion, collagenase type II (255 U/mg, Worthington Biochemical) was introduced and the heart was perfused for an additional 15 minutes until the heart became soft. The ventricles were minced and digested in Krebs-Henseleit buffer containing 1% BSA. Calcium then was reintroduced slowly to cardiac myocytes at a final concentration of 1.25mM. Cells were counted, plated on laminin (Sigma) coated-plates for calcium imaging or plated in 48-well plate at an equal cell density (2^10^4^ cells) for cell viability studies. Cells were maintained in media 199 (Invitrogen) containing 1% BSA and placed in a 37° incubator for use the next day.

The next day, to determine the impact of acetaldehyde on calcium influx, adult cardiac myocytes were incubated for 40 min at 37°C in DMEM serum free medium (Invitrogen). Cover slips containing cardiac myocytes were mounted on a Zeiss Axiovert inverted fluorescent microscope. Cardiac myocytes were superfused continuously with DMEM serum free media by a peristaltic pump at a flow rate of 2 mL/min. The basal intracellular calcium activity was recorded for 5 minutes. Cells were then superfused with a pulse of acetaldehyde (0.1, and 1 μM) in DMEM serum free media for 30 seconds to mimic an exposure occurring when inhaling an e-cigarette. This was followed by KCl (60mM) as a positive control. Images and ratio metric real-time calcium tracing data were acquired using an alternating excitation wavelength (340 and 380 nm) and emission wavelength (510 nm). Background fluorescence was corrected by Easy Ratio Pro software. The ratio of the 2 intensities acquired by Easy Ratio were used to measure changes in intracellular calcium levels, as previously described [24].

In a separate study, wild type ALDH2 and ALDH2*2 knock-in primary cardiac myocytes were incubated with acetaldehyde (0.1 and 1 μM) for 4 hours and cell viability was measured by 3-(4,5-dimethylthiazol-2-yl)-2,5-diphenyltetrazolium bromide (MTT) assay. The yellow tetrazolium dye MTT was reduced to purple formazan in living cells. The absorbance of formazan was quantified at 560 nm using a Synergy 2 plate reader (BioTek, Winooski, Vermont, United States).

### 2.5. ROS activity assay

Oxidative stress was measured by detecting intracellular ROS accumulation in H2-DCFDA (2’,7’-dichlorofluorescein diacetate) assay [25]. Briefly, the H2-DCFDA was deacetylated by a viable cell to a non-fluorescent compound, which was further oxidized by reactive oxygen species (ROS) into a highly fluorescent compound DCF. The wild type ALDH2 and ALDH2*2 knock-in primary cardiac myocytes were incubated 10 μM H2-DCFDA in medium at 37 °C for 30 min, and then the medium was removed and washed with PBS. Next cells were incubated with medium, medium, medium with 0.1 or 1 μM acetaldehyde or 20 μM 4-HNE as positive control. DCF fluorescence was measured for 2 hours at excitation and emission wavelengths of 485 and 535 nm, respectively.

### 2.6. Immunohistochemistry and confocal microscopy

Primary cardiomyocytes from both wild type ALDH2 mice and ALDH2*2 mice were seeded on laminin precoated coverslip glass for immunofluorescence. Cardiomyocytes were preincubated with M199 medium with vehicle or acetaldehyde 1 μM for 30 minutes. Cardiomyocytes were fixed with 4% paraformaldehyde for 10 minutes at room temperature. Next, cardiomyocytes were incubated with primary antibodies (anti-dystrophin 1:200, Abcam; anti-4HNE 1:200, Alpha Diagnostic) overnight at 4°C, then incubated with fluorophore-conjugated secondary antibodies the next day (Alexa Fluor 488 or Alexa Fluor 594, Invitrogen) at 1:1000 dilution. Cardiomyocytes were mounted to glass slides using anti-fade mountant with DAPI (Thermofisher Scientific) and imaged using a Zeiss LM900 confocal microscope.

### 2.7. Telemeter implantation for EKG recording

To monitor heart rate with e-cigarette exposures in conscious freely moving rodents, male mice were implanted with wireless telemetry systems to record the electrocardiogram (KAHA Sciences, Grafton, New Zealand). Mice were placed in the supine position on a heating pad to maintain the body temperature. Telemeters were implanted posteriorly under isoflurane anesthesia with EKG leads tunneled and placed in a lead II configuration. The surgical incision was closed with non-absorbable 4-0 veterinary sutures and observed for any infection or necrosis. After surgery, mice were given one week to recover. Animal were monitored continuously until able to ambulate freely, and post-operative analgesic was given (Buprenorphine, subcutaneous injection, 0.1 mg/kg) if severe pain was observed. Additionally, 4 days prior to e-cigarette exposure, telemeter functionality was tested by administering labetolol (10 mg/kg, intraperitoneal injection) to rodents while measuring heart rate from the remote telemeter captured by placing rodents on a wireless digital receiver tBase (Model: MT110, KAHA Sciences) with data continuously steamed and recorded by LabChart (AD Instruments, Sydney, Australia). After confirmation of the telemeter functionality, heart rate was measured while rodents were exposed to room air or e-cigarette aerosol.

### 2.8. E-cigarette aerosol exposure and tissue collection

After telemeter implantation and testing, rodents were divided into 4 groups: room air exposed wild type ALDH2 (WT) mice, room air exposed homozygous ALDH2*2 mice, e-cigarette exposed wild type mice, and e-cigarette exposed ALDH2*2 mice. The mice were exposed to e-cigarette aerosol within a 2L exposure chamber, where the aerosol to air ratio was 1:6. To monitor gas levels within the chamber, a multi-gas monitor (BW Honeywell, Charlotte, NC) continuously measured oxygen and carbon monoxide concentrations and was set to alarm if the environmental oxygen levels dropped by 1%.

The morning before exposure, the baseline heart rate for each of the four groups was recorded for 5 minutes. Exposures were then performed by pairing a wild type ALDH2 mouse with an ALDH2*2 mouse. Aerosol from a JUUL e-cigarette or room air was drawn and delivered to the exposure chamber by a peristaltic pump with a 20mL total puff volume per exposure. Mice were exposed to e-cigarette aerosol or room air for 4 sessions a day for 10 days. Next, for each exposure session, 7 puffs/min were drawn for the first two minutes (a total of 14 puffs), and the animal whole body exposure to JUUL aerosol continued for an additional 5 minutes (exposure phase) followed by 23 minutes of a smoke-free interval for each session (recovery phase). Heart rate was recorded daily. The puff volume and exposure duration were based on prior studies [26–29].

After completing 10 days of exposure, rodents were anesthetized with 100 mg/kg Inactin (100mg/ml intraperitoneal, Sigma-Aldrich) and euthanasia were confirmed with failure of responding to toe pinch. The hearts were removed and homogenized in mannitol sucrose buffer (210 mM mannitol, 70 mM sucrose, 1 mM EDTA, 5 mM MOPS) and centrifuged at 10000 rpm for 10 minutes at 4°C to remove the cell debris. The supernatants were transferred to 1.5 ml Eppendorf tube and placed at −80° freezer for further molecular studies.

### 2.9. Protein carbonylation

Carbonyl groups introduced into protein side chains (reflecting aldehyde-induced oxidative modification of proteins) were measured by an OxyBlot protein oxidation detection kit (Millipore) by equally splitting the heart homogenate into two samples. One sample was derivatized by adding DNPH (2,4-dinitrophenylhydrazine) solution. The carbonyl groups in the protein side chains were derivatized to 2,4-dinitrophenylhydrazone (DNP-hydrazone) by reaction with DNPH. As a negative control, the other sample was mixed with derivatization-control solution. Specifically, 15-20 μg of protein was added into 10 μL of the positive/negative derivatization reaction solution, and the mixtures were incubated at room temperature for 15 minutes. Next, the DNP-derivatized protein were separated and detected by western blot.

### 2.10. Western blot detection for 4HNE protein adducts, protein carbonylation and NF-κB

The formation of 4-hydroxynonenal-induced protein adducts, carbonylated protein, and nuclear factor kappa B (NF-κB/p65) in heart homogenates were analyzed by western blot. The protein content was determined by Pierce BCA protein assay kit (Thermo Scientific). Samples were resolved on 10% SDS-polyacrolamide gels at 200 volts for 2 hours at room temperature and transferred to PVDF membranes at 100 volts for 90 minutes at 4°C. The membranes were then incubated with primary antibody (anti-4HNE from Millipore 1:1000; anti-DNP 1:500; anti-phospho-NFκB from Invitrogen 1:1000; anti-NFκB from Invitrogen 1:1000) at 4°C overnight, followed by incubation with secondary antibody (anti-rabbit from Invitrogen 1:2000) at room temperature for 2 hours. Membranes were incubated for chemiluminescence at room temperature for 5 minutes and imaged by using an Azure cSeries gel imaging system (Azure Biosystems, Dublin, CA).

### 2.11. Free MDA production

Free malondialdehyde (MDA) was measured by reacting with thiobarbituric acid (TBAR) to generate a MDA-TBAR adduct according to the manufacturer’s instructions (Abcam, Cambridge, MA). Briefly, the heart homogenates were lysed in manufacturer provided lysis buffer with butylated hydroxytoluene (BHT) and centrifuged at 13,000g for 10 minutes to collect the supernatants. For each well containing MDA standard and samples, 3x volumes of TBA reagent was added to generate the MDA-TBA adduct. The absorbance of MDA-TBA was measured at 532nm using a Synergy 2 plate reader (BioTek, Winooski, Vermont, United States).

### 2.12. Statistics

Based upon power analysis, a minimum of 6 animals was necessary to achieve at least a 20% minimal difference in heart rate changes in e-cigarette exposed mice for a power of 95% with α<0.05 and β<20%. Data analysis were performed using Graph Pad Prism 7.0 software. Data were expressed as mean ± S.E.M. The *p* value was calculated by ANOVA with Bonferroni correction was performed for multiple comparisons between groups and Student’s t-test was performed when comparing two treatments, with corresponding number (n) indicated in figure legends. *^#+^*p*<0.05.

## 3. Results

### 3.1 Quantification of nicotine and aldehydes in e-cigarette aerosol

Three e-cigarette brands (Blu, Halo and JUUL) were quantified for nicotine and aldehydes (Figure 1A). Aerosolized nicotine for Blu (24 mg/ml), Halo (24 mg/ml), and JUUL (50 mg/ml) e-cigarettes were measured relative to air (Figure 1B, Blu: 0.6±4.2* ppm, Halo: 9.9±4.1* ppm, JUUL: 22.6±2.0* ppm versus air: 0.001±0.0001 ppm, respectively, n=20/group, \**p*<0.01 versus air). In addition, acetaldehyde levels were highest in the e-cigarette Blu, followed by Halo and JUUL, relative to air (Figure 1C, Blu: 10.7±4.6* ppm, Halo: 8.6±3.4* ppm, JUUL: 5.3±1.4* ppm versus air: 0.004±0.0002 ppm, respectively, n=20/group, \**p*<0.01 versus air). Formaldehyde levels in e-cigarette aerosols were several-fold less relative to acetaldehyde (Figure 1D, Blu: 0.8±0.6* ppm, Halo: 0.7±0.5* ppm, JUUL: 0.2±0.08* ppm, versus air: 0.005±0.0004 ppm, respectively, n=20/group, \**p*<0.01 versus air). Levels of acrolein were also substantially lower than acetaldehyde (Figure 1E, Blu: 1.2±0.9* ppm, Halo: 0.6±0.4* ppm, JUUL: 0.09±0.06* ppm, versus air: 0.002±0.0004 ppm, respectively, n=20/group, \**p*<0.01 versus air). Additionally, the total aldehyde load (acetaldehyde, formaldehyde, acrolein) varied between the 3 e-cigarette brands and linearly correlated with the ratio of propylene glycol to vegetable glycerin (Supplemental Figure 1, JUUL (30/70): 5.6±0.3 ppm, Halo (50/50): 9.9±0.9 ppm, Blu (60/40): 12.6±1.3 ppm, r= 0.99, p<0.0025). Together, these results identify that e-cigarette aerosol primarily contains acetaldehyde with other aldehydes several-fold less within e-cigarette aerosol.

**Figure 1.**
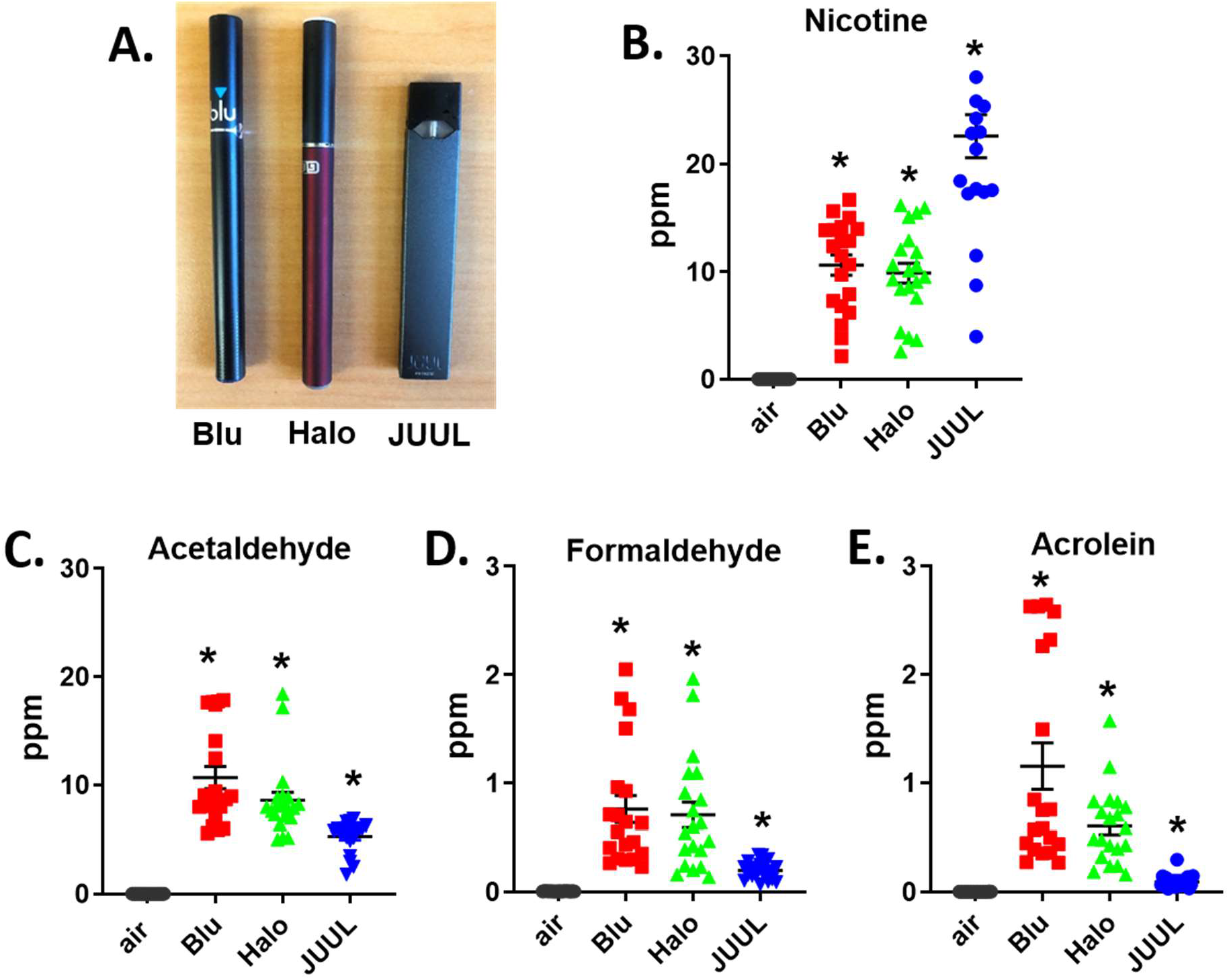
E-cigarette aerosol nicotine and aldehyde quantification. **A.** Three brands of e-cigarettes. Amount of **B.** nicotine **C**. acetaldehyde **D.** formaldehyde and **E**. acrolein present in e-cigarette aerosol or room air. Data was expressed as mean ± SEM (ppm). \**p*<0.05 Blu, Halo and JUUL versus air, calculated by Student’s t-test. n= 20 measurements per e-cigarette.

### 3.2. Basal ALDH2 expression and activity, ROS, and nuclear factor kappa B (NF-κB)

As e-cigarettes primarily contained acetaldehyde and the enzyme responsible for metabolizing acetaldehyde is ALDH2, we used wild type ALDH2 and ALDH2*2 knock-in rodents (Figure 2A). We quantified ALDH2 protein expression and activity for wild type ALDH2 and ALDH2*2 mice in heart, lung, and liver homogenates. Interestingly, the relative protein expression in heart homogenates for wild type ALDH2 mice is 2-fold lower relative to the lung and liver by Western blot. (Figure 2B, heart 0.45±0.02^#^ relative to lung 1.06±0.03 and liver 1.27±0.04, ALDH2/GAPDH relative densitometry units, respectively, n=3/group, #p<0.05 versus lung and liver). As expected, the protein expression for ALDH2*2 rodents is also 2 to 3-fold lower compared to wild type ALDH2 rodents. For the ALDH2*2 rodents there is also lower ALDH2 protein expression for the heart relative to the lung and liver (Figure 2B, heart 0.24±0.01^#^ relative to lung 0.58±0.01 and liver 0.60±0.02, ALDH2/GAPDH relative densitometry units, n=3/group, #p<0.05 versus lung and liver). The expression of ALDH2 for either rodent was similar between genders (Supplemental Figure 2). The ALDH enzymatic activity for the wild type ALDH2 heart is also 1.5-fold lower relative to the wild type ALDH2 lung and liver (Figure 2C, heart 4.7±0.88 relative to lung 6.7±1.52 and liver 7.2±1.36, μmol/min/mg protein, n=3/group, #*p*<0.05 versus heart). The tissue type, protein expression, and point mutation within an integral α-helix of the enzyme causes ALDH activity for the ALDH2*2 heart homogenate to have 2 to 3-fold lower activity when comparing to the respective wild type ALDH2 organ. The ALDH2*2 heart has less ALDH activity relative to the ALDH2*2 lung and ALDH2*2 liver (Figure. 2C, heart 1.3±0.25^#^ versus lung 2.1±0.17 and liver 2.7±0.26, μmol/min/mg protein, n=3, #p<0.05 versus heart, *p<0.05 ALDH2*2 versus wild type ALDH2). Further, ALDH2*2 primary adult cardiomyocytes demonstrated 2-fold higher levels of intracellular ROS relative to ALDH2 wild type cardiomyocytes (Figure 2D. RFU: ALDH2*2: 22678±4354* versus ALDH2: 10859 ± 2057, n=4/group, *p<0.05 between ALDH2*2 mice and wild type mice). ROS overproduction is also known to activate NF-κB by phosphorylation [30] and for ALDH2*2 heart homogenates p-NF-κB was elevated 2-fold relative to wild type ALDH2 hearts (Fig. 2E, ALDH2*2: 0.42±0.03* versus WT: 0.25±0.01, respectively, ratio of p-NF-κB to total NF-κB, n=3/group, *p<0.05 between ALDH2*2 mice and ALDH2 wild type mice).

**Figure 2.**
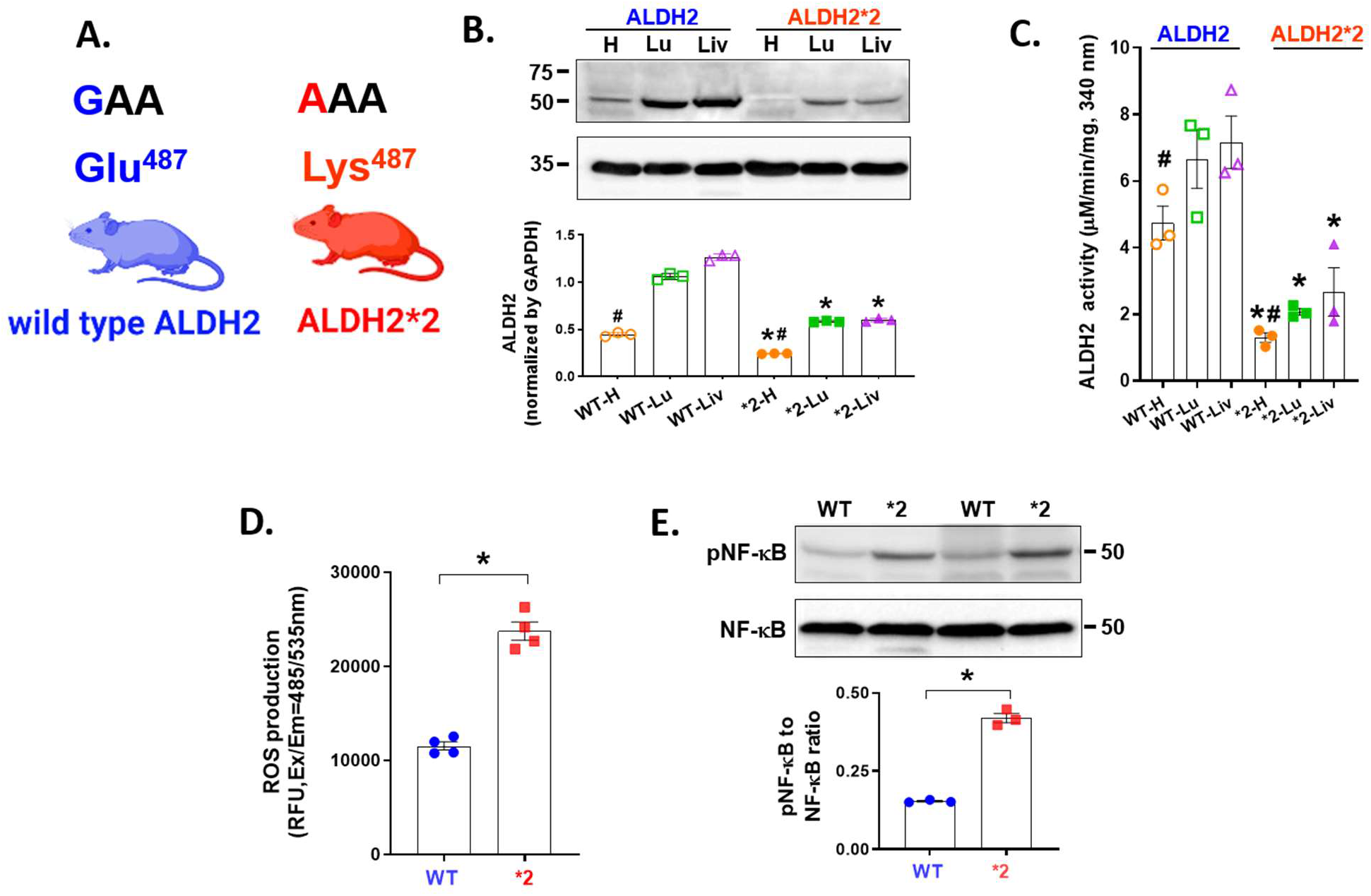
Basal levels of ALDH2 expression and activity, ROS and nuclear factor kappa B (NF-κB). **A**. An ALDH2 knock-in rodent on a C57/BL6 background was generated with a missense mutation (glutamic acid to lysine at 487) that reflects the human ALDH2*2 genetic variant. **B**. Top: Representative western blot of ALDH2 (56 kDa) protein expression for heart, lung, and liver tissue homogenates for ALDH2 and ALDH2*2 mice. Bottom: Quantification of ALDH2 protein expression normalized to GAPDH (36 kDa, n=3/group). **C.** ALDH2 enzymatic activity for homogenates of each genotype and organ (n=3/group). **D.** ROS levels in cardiomyocytes from wild type ALDH2 and ALDH2*2 mice (n=4/group). **E**: phospho- and total nuclear factor kappa B (NF-κB, 50 kDa) in heart homogenates from unexposed wild type ALDH2 and ALDH2*2 mice with representative western blot of phospho- and total NF-κB expression and quantification (n=3/group). Blue represents wild type ALDH2 mice and red represents the ALDH2*2 mice. WT= wild type, *2 = ALDH2*2, H= heart, Lu= lung, Liv= liver. \**p*<0.05 comparison between wild type ALDH2 mice and ALDH2*2 mice, #*p*<0.05 hearts relative to other organs of the same genotype, calculated by Student’s t-test.

### 3.3. primary isolated cardiomyocytes response to acetaldehyde incubation

When challenging ALDH2 and ALDH2*2 primary adult cardiomyocytes with an acetaldehyde pulse to mimic an e-cigarette, a clear difference in the dose-dependent calcium influx occurred between the genotypes. ALDH2*2 myocytes responded with an intracellular calcium influx even at the lowest acetaldehyde dose tested (0.1 μM), as opposed to wild type ALDH2 myocytes (Figure 3A: 6.7±0.6 versus 0.7±0.3, respectively, % change relative to maximal peak response, n =21 cells for ALDH2 and n=20 cells for ALDH2*2 from 4 biological replicates, p<0.05 ALDH2*2 cardiomyocytes versus wild type ALDH2 cardiomyocytes). The difference in calcium influx between the ALDH2 genotypes continued to separate dose-dependently. At a 10-fold higher dose of acetaldehyde (1 μM), there was a 4-fold greater peak response (Supplemental Figure 3: 19.1±5.2 versus 4.0±0.6, respectively, % maximal peak response, n=19 cells for ALDH2 and n=17 cells for ALDH2*2 from 4 biological replicates, p<0.05 ALDH2*2 versus wild type ALDH2). The peak maximal response triggered by 0.1-1 μM acetaldehyde is summarized (Figure 3B). In the presence of acetaldehyde, ALDH2*2 myocytes had a higher intracellular ROS level relative to wild type ALDH2 myocytes (Figure 3C: 0.1 μM acetaldehyde, RFU, ALDH2*2: 34106±1488^*^ versus ALDH2: 14437.0±1010, respectively, \**p*<0.05 between wild type ALDH2 mice and ALDH2*2 mice. ^+^*p*<0.05 comparison between acetaldehyde treated cardiac myocytes relative to its vehicle control group). 1 μM acetaldehyde, RFU, ALDH2*2: 42756±798 versus ALDH2: 19323±1237, respectively, n=4/group, * *p*<0.05 ALDH2*2 versus wild type ALDH2).

**Figure 3.**
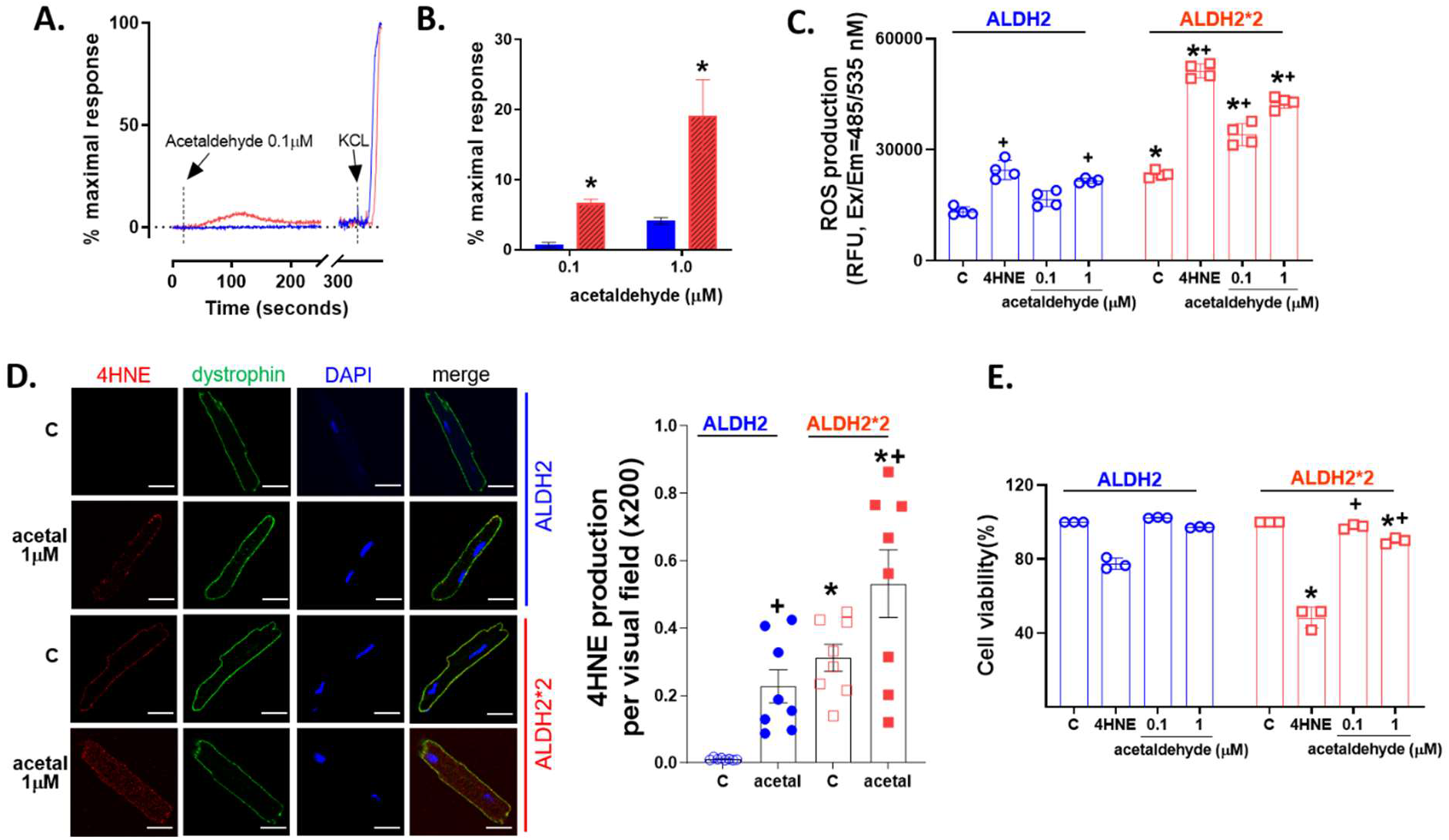
Cardiac myocyte response to acetaldehyde in rodents. **A.** Calcium influx by 0.1 μM acetaldehyde. **B**. Max peak response triggered by 0.1 or 1μM acetaldehyde relative to 60mM KCl. (n=21 cells ALDH2 and n=20 cells ALDH2*2 from 4 biological replicates). **C.** ROS activity by 0.1 or 1μM acetaldehyde with 4-HNE (20μ as a positive control (n=4 biological replicates/treatment). **D**. Left: Immunostaining of primary cardiomyocytes for 4-hydroxynoneal (4-HNE) protein adducts in vehicle or acetaldehyde treated cells, scale 20 μm. Green: dystrophin, Red: 4-HNE protein adducts, Blue: DAPI. Right: Quantification of 4-HNE protein adducts (n=8/group). **E.** Acetaldehyde-induced cell death as a percentage of cell viability compared with vehicle group. n= 3 biological replicates/treatment. \**p*<0.05 comparison between wild type ALDH2 mice and ALDH2*2 mice, ^+^*p*<0.05 comparison between acetaldehyde treated cardiac myocytes relative to its vehicle control group, calculated by Student’s t-test.

ALDH2*2 primary cardiomyocytes also demonstrated higher baseline levels of 4-HNE protein adducts relative to wild type ALDH2 cardiomyocytes, which was increased by application of 1 μM acetaldehyde (Figure. 3D: Untreated, ALDH2*2: 0.31±0.11* compared with ALDH2: 0.01±0.01; 1 μM acetaldehyde treated, ALDH2*2: 0.53±0.28* compared with wild type ALDH2: 0.31±0.11, n=8 figures taken from 3 slides of cardiomyocytes isolated from 3 mice in each treatment group, \**p*<0.05 between wild type ALDH2 mice and ALDH2*2 mice. ^+^*p*<0.05 comparison between acetaldehyde treated cardiac myocytes relative to its vehicle control group). Further, the lowest dose of acetaldehyde tested had no effect on cardiac myocyte cell viability for the wild type ALDH2 or the ALDH2*2 cardiac myocyte (Fig 3E, 0.1 μM: 102±3% versus 98±1%, respectively, n=3/group). Acetaldehyde levels at the higher dose induced cell death which was greater for ALDH2*2 cardiac myocytes relative to wild type ALDH2 cardiac myocytes (Fig. 3E: 1 μM: 97±4% versus 90±2%*, respectively, n=3/group). *p<0.05 ALDH2*2 versus wild type ALDH2. #p<0.05 relative to untreated control.

### 3.4. E-cigarette exposure impact on heart rate in rodents

As acetaldehyde is highly prevalent in all e-cigarette aerosols tested, we questioned what impact the inactivating genetic variant, ALDH2*2 relative to wild type ALDH2, has on e-cigarette aerosol exposure in rodents. We first quantified heart rate physiologically during and after exposure (as acetaldehyde accumulation after alcohol consumption results in tachycardia for humans carrying the ALDH2*2 variant) [31, 32]. We implanted 38 male rodents (18 wild type and 20 ALDH2*2) with telemeters to continuously monitor heart rate in conscious and freely moving rodents. After telemeter instrumentation, 4 mice (1 wild type and 3 ALDH2*2 mice) were excluded prior to entering a protocol due to wound dehiscence at the site of the implanted telemeter. A representative EKG waveform in normal sinus rhythm from our mouse model (Supplemental Fig. 4A). To confirm the telemeters can identify real-time heart rate changes, we administered an intraperitoneal injection of labetolol or saline. During this process, 2 mice (1 wild type and 1 ALDH2*2) died due to intravascular injection and did not make it into the study. Labetolol given to wild type ALDH2 and ALDH2*2 mice decreased heart rate relative to saline treated 1 minute after injection (Supplemental Figure 4B, labetolol treated: wild type 538±108 bpm, ALDH2*2 467±145 bpm relative to saline wild type: 697±61 bpm, ALDH2*2: 721±38 bpm, respectively, n=8/group, *p<0.0001 relative to saline treated).

Next, wild type ALDH2 and ALDH2*2 mice were exposed to e-cigarette aerosol or air four times per day for 10 days (Figure 4A). During exposures, mice were exposed paired by genotype to e-cigarette aerosol or air while continuously monitoring oxygen and carbon monoxide levels within the exposure chamber (Figure 4B). Aerosol from an e-cigarette or room air was drawn and delivered to the exposure chamber by a peristaltic pump. Using real-time mass spectrometry, we also quantified the concentrations of nicotine, acetaldehyde, formaldehyde, and acrolein for e-cigarette aerosol or air within the exposure chamber (Figure 4C, nicotine 12±11* ppm versus air 0.03±0.02 ppm; 4D: acetaldehyde 0.8±0.4* ppm versus air 0.03±0.01 ppm; 4E: formaldehyde 0.04±0.01* ppm versus air 0.01±0.007 ppm, and 4F: acrolein 0.008±0.002* ppm versus air 0.005±0.003 ppm, respectively, e-cigarette aerosol versus air, n=20/group, *p<0.05).

**Figure 4.**
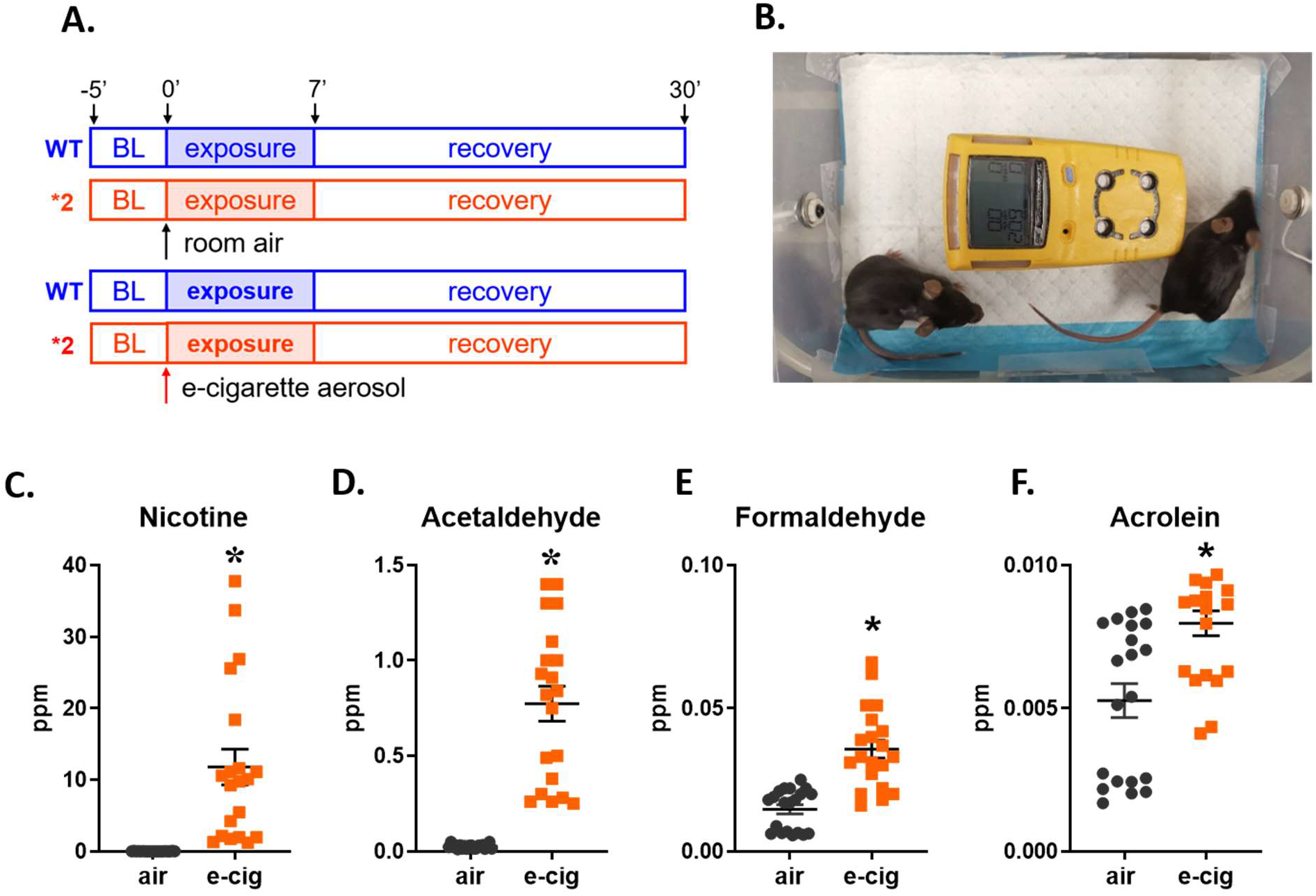
E-cigarette aerosol exposure protocol and quantification of aerosol within the exposure chamber. **A**. Experimental protocol. Wild type ALDH2 mice or ALDH2*2 mice were exposed to 7 minutes of e-cigarette aerosol exposure four times daily for 10 total days. **B**. Picture of the e-cigarette aerosol exposure chamber, where rodents were pair-matched (one wild type ALDH2 and one ALDH2*2 rodent) to an exposure. Device in yellow is a multi-gas monitor. **C-F**. E-cigarette aerosol average concentration of nicotine and aldehydes in the rodent exposure chamber. **C.** nicotine; **D.** acetaldehyde; **E.** formaldehyde; and **F.** acrolein, \**p*<0.05 comparison between e-cigarette aerosol and air, calculated by Student’s t-test.

At baseline, there is no significant difference in the resting heart rate occurring between rodent groups (Figure 5A: air exposed: ALDH2 mice 525±61 bpm and ALDH2*2 mice 543±42 bpm versus e-cigarette aerosol: ALDH2 mice 516±55 bpm and ALDH2*2 mice 525±42 bpm, respectively, n=8 per group). An increase in heart rate was evident midway through the exposure protocol in the ALDH2*2 mice unlike the wild type ALDH2 mice (Fig 5B, air exposed: ALDH2 mice 474±48 bpm and ALDH2*2 mice 486±85 bpm versus e-cigarette aerosol: ALDH2 mice 546±58* bpm and ALDH2*2 mice 637±85* bpm, respectively, n=7-8/group, *p<0.05 versus the air exposed mice). Exposure to e-cigarette aerosol over 10 days caused the baseline resting heart rate to increase for rodents exposed to e-cigarettes. This was unlike the air exposed groups (Fig. 5B, ALDH2 mice: 462±99 to 512±64 bpm, ALDH2*2 mice: 506±58 ppm to 530±75 bpm, n=7-8/group). Additionally, after completing the daily e-cigarette exposure, the e-cigarette aerosol exposed ALDH2*2 mice started to demonstrate an increased baseline heart rate starting from midway through the exposure protocol relative to the heart rate measured for the first day after exposure (Fig 5C: ALDH2: 557±90 versus 601±62* bpm, ALDH2*2: 568±98 versus 678±61* bpm, n=7-8/group, *p<0.05 versus the air exposed mice). On the final day after exposure, e-cigarette aerosol exposed wild type ALDH2 and ALDH2*2 mice both demonstrated higher heart rates relative to the air exposed mice (Fig. 3C: e-cigarette aerosol: ALDH2 mice 646±49* bpm and ALDH2*2 mice 670±132* bpm, n=7-8/group, *p<0.05 versus the air exposed mice). These increases in heart rate were not seen for the air-exposed wild type ALDH2 or ALDH2*2 mice (Fig 5C).

**Figure 5.**
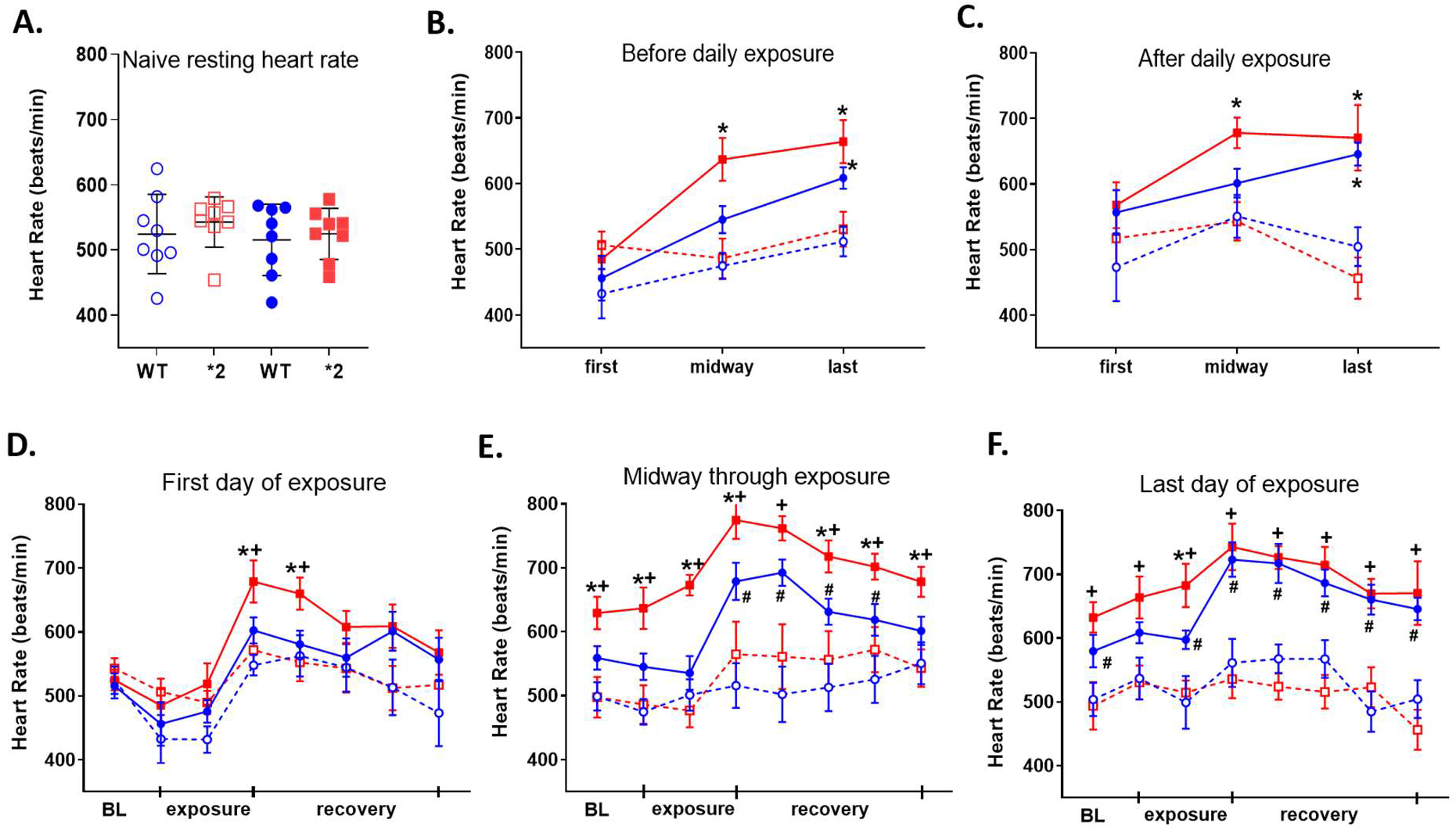
Heart rate measurements of ALDH2 and ALDH2*2 mice exposed to e-cigarette aerosol. **A.** Baseline heart rate before exposure. **B**. Mean heart rate prior to exposure over the 10-day course. **C**. Mean heart rate after the completion of daily exposure over the 10-day course. **D**. Mean heart rate changes during the first exposure **E**. midway through the 10-day exposure and **F**. the last day of the 10-day exposure. Data is expressed as mean ± SEM. Solid colors are rodents exposed to e-cigarette aerosol, dashed line and open circles are rodents exposure to air. Blue is wild type ALDH2 and red is ALDH2*2 rodents. \**p*<0.05 comparison between e-cigarette exposed ALDH2*2 mice and ALDH2 wild type mice, **+***p*<0.05 comparison between e-cigarette exposed ALDH2*2 mice and room air exposed ALDH2*2 mice, #*p*<0.05 comparison between e-cigarette exposed wild type ALDH2 mice and air exposed wild type ALDH2 mice, calculated by two-way ANOVA with Bonferroni correction. n=7-8/group.

Further, with exposure to e-cigarette aerosol or air, the heart rate peaked at the end of e-cigarette aerosol exposure continuing into the recovery phase. During the first day of e-cigarette aerosol or air exposure, only ALDH2*2 mice showed a significantly elevated heart rate after e-cigarette aerosol relative to air exposure (Fig. 5D: 679±93*^+^ bpm versus 572±81 bpm, respectively, n=8 for Juul-WT and air-ALDH2*2, n=7 for air-WT and Juul-ALDH2*2, *^+^p<0.05). In contrast, wild type ALDH2 mice exposed to e-cigarette aerosol showed a trend of increasing heart rate which was not significant in e-cigarette aerosol relative to air exposed rodents (Figure 5D, 603±54 bpm versus 573±61 bpm, respectively, n=8 for Juul-WT and air-ALDH2*2, n=7 for air-WT and Juul-ALDH2*2). Midway through the 10-day exposure period, both wild type ALDH2 and ALDH2*2 mice exhibit a significant increase in heart rate when exposed to e-cigarette aerosol relative to air (Fig. 5E: e-cigarette aerosol ALDH2*2 mice 775±30*^+^ bpm and ALDH2 679±82^#^ bpm, air exposure ALDH2*2 mice 565±144 bpm mice and ALDH2 516±93 bpm, n=7-8/group, *^+#^ p<0.05). During the 10^th^ day of e-cigarette aerosol or air exposure, the ALDH2*2 and wild type ALDH2 mice subjected to e-cigarette aerosol had increased heart rates relative to rodents exposed to air (Fig. 5F, e-cigarette aerosol: ALDH2*2 mice 743±96^+^ bpm and wild type ALDH2 mice 723±77^#^ bpm versus ALDH2*2 mice 536±85 bpm and wild type ALDH2 561±106 bpm, n=7-8/group, ^+#^ p<0.05).

### 3.5. E-cigarette aerosol exposure causes cardiovascular oxidative stress in rodent hearts

After e-cigarette aerosol or room air exposure, we quantified cardiovascular oxidative stress by measuring protein carbonylation, total free malondialdehyde (MDA) production, and 4-hydroxynonenal (4HNE)-protein adduct formation in heart homogenates after the 10 days of e-cigarette aerosol or room air exposure.

Protein carbonylation was quantified by western blot (Fig. 6A, Supplemental Figure 5). The protein carbonylation levels were elevated when ALDH2*2 mice or wild type mice were exposed to e-cigarette aerosol compared to air exposed mice (Fig. 6B, e-cigarette: ALDH2*2 mice 119±3*^#^ and ALDH2 mice 95±1* relative to air: ALDH2*2 mice 79±2 and ALDH2 mice 54±2, respectively, ratio of protein carbonylation to negative control, n=3/group, *p<0.05 between ALDH2*2 mice and wild type ALDH2 mice, +p<0.05 between e-cigarette exposed mice and air exposed mice). Free MDA was also elevated in heart homogenates from ALDH2*2 mice 2-fold relative to homogenates from air exposed ALDH2*2 mice (Fig.6C, e-cigarette: ALDH2*2 mice 99±13* vs ALDH2 mice 59±7, nmol/ml MDA, n=3/group, *^#^ p<0.05). This was in contrast to wild type ALDH2 heart homogenates which did not show increased levels of free MDA relative to air treated wild type mice homogenates (Fig. 6C, 55±7 vs 43±11, nmol/ml MDA, n=6/group). Further 4-HNE-induced protein adducts were elevated 1.5-fold for the ALDH2*2 mice in the e-cigarette exposed group relative to the air exposed group (Figure. 6D, 58±2^+^ vs 38±6, n=3/group, ^#^p<0.05). This elevation of 4-HNE-induced protein adducts was also 1.5-fold higher in wild type ALDH2 mice exposed to e-cigarette aerosol relative to air (Figure 6E, 57±2 vs 38±2, n=3/group, ^#^p<0.05). Heart homogenates from both genotype mice subjected to e-cigarette aerosol had a significant increase in p-NF-kB relative to homogenates from rodents exposed to air (Fig. 6F, e-cigarette aerosol: ALDH2*2 mice 0.68±0.03 and wild type ALDH2 mice 0.51±0.04 versus air: ALDH2*2 mice 0.42±0.03 and wild type ALDH2 mice 0.25±0.01, n=3/group, p-NF-κB to NF-κB ratio, *p<0.05 between ALDH2*2 mice and wild type mice, ^+^p<0.05 between e-cigarette aerosol exposed mice and air exposed mice).

**Figure 6.**
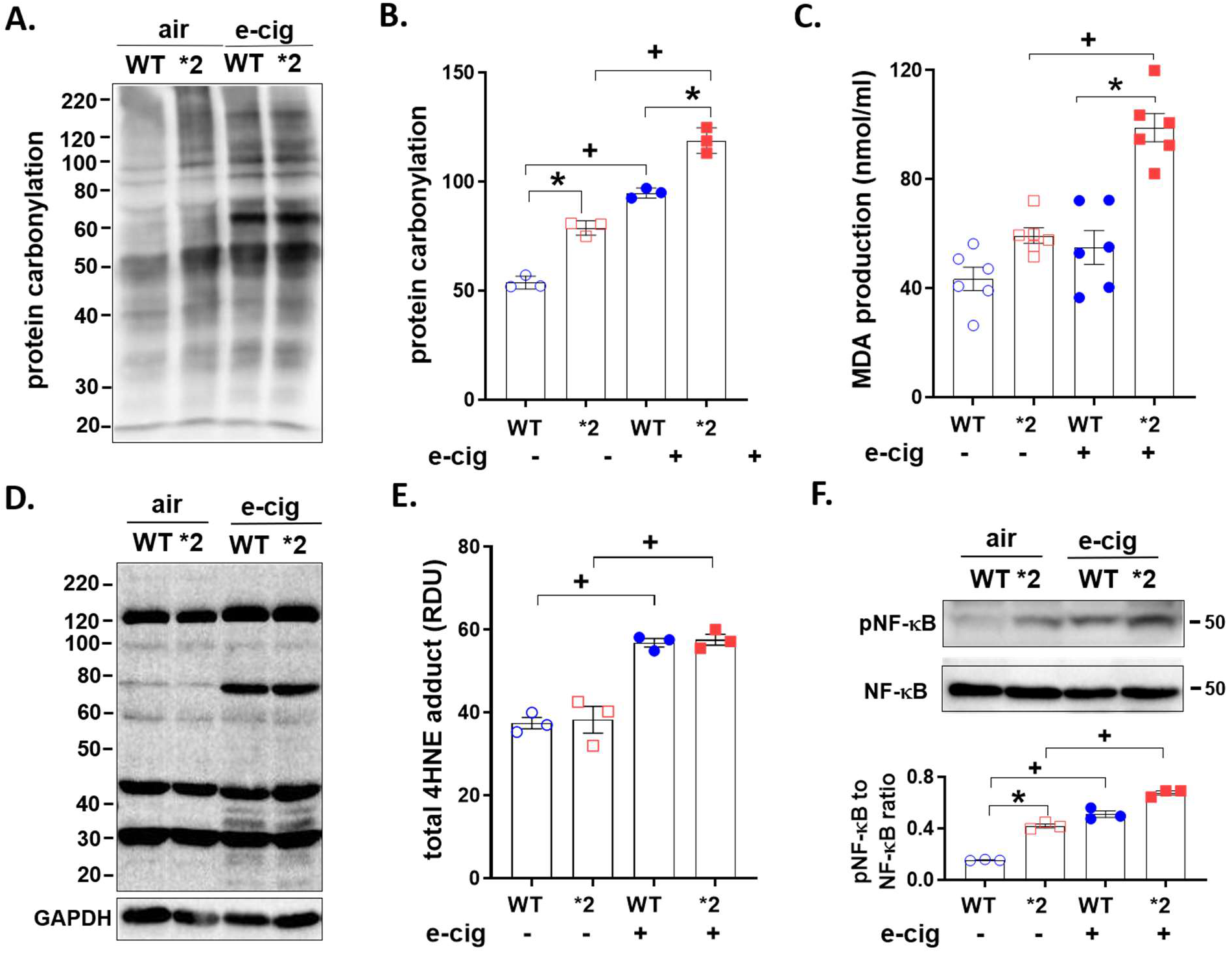
Aldehydic load after e-cigarette aerosol or air exposure. **A**. Representative blot of protein carbonyl expression. **B**. Quantification of protein carbonylation (n=3/group). **C.** Free malondialdehyde (MDA) in heart homogenates. The concentration of MDA was normalized by the total amount of protein (n=6/group). **D**. Representative blot of 4-HNE-induced protein adducts with GAPDH (36 kDa) as a loading control. **E.** Quantification of 4-HNE-induced protein adducts (n=3/group). **F**: phospho- and total nuclear factor kappa B (NF-κB, 50kDa) in heart homogenates (n=3/ group). \**p*<0.05 comparison between wild type ALDH2 mice and ALDH2*2 mice, ^+^*p*<0.05 comparison between e-cigarette exposed mice and room air exposed mice, calculated by Student’s t-test.

## 4. Discussion

Here we identified acetaldehyde as a primary aldehyde within e-cigarette aerosol. When exposed to e-cigarette aerosol, the ALDH2*2 mice (due to genetics that limit acetaldehyde metabolism) are more susceptible to cardiovascular oxidative stress when compared to wild type ALDH2 rodents. As the ALDH2*2 mice are a knock-in model where a single copy of the ALDH2 gene is replaced with the inactive human ALDH2*2 variant, this study provides valuable insight that people carrying an ALDH2*2 variant may potentially be more susceptible to elevated cardiovascular oxidative stress that can be elevated with frequent e-cigarette aerosol exposure. Exposure to aldehydes within e-cigarette aerosol can promote ROS production, increased protein carbonylation, lipid peroxidation, and NF-κB activation leading to elevated levels of cardiovascular oxidative stress that is further exacerbated in the ALDH2*2 variant due to the limited ability to metabolize aldehydes within e-cigarette aerosol (Figure 7). This may pave the way to developing precision medicine strategies towards understanding cardiovascular risk for e-cigarette exposures when considering genetics influencing aldehyde metabolism.

**Figure 7.**
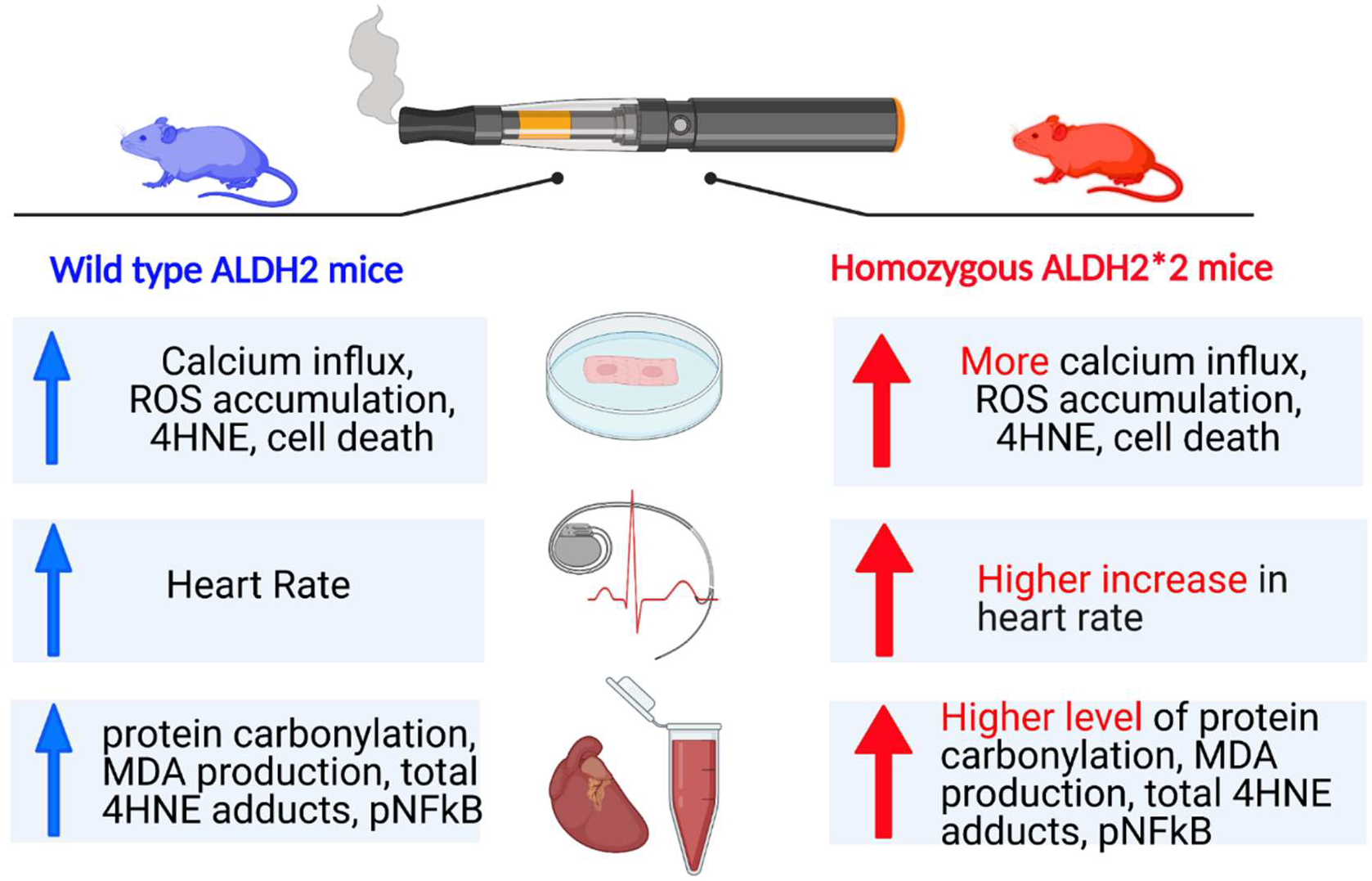
Summary Figure. Graphical summary of differences for wild type ALDH2 mice and ALDH2*2 mice when exposed to e-cigarette aerosol.

Aldehydes are generated from e-cigarettes by the incomplete combustion of propylene glycol and vegetable glycerin when exposed to oxygen [33]. Although e-cigarette aerosol contains aldehydes, the levels could be as much as 70-fold less compared to conventional cigarette smoke [2]. Our findings identify within e-cigarette aerosol aldehydes including acrolein, acetaldehyde, and formaldehyde consistent with prior studies [2, 34–36]. By using real-time mass spectrometry to measure aldehydes, our study identifies acetaldehyde as the primary aldehyde within e-cigarette aerosol. This is consistent with a prior study identifying acetaldehyde as the predominant aldehyde within e-cigarette aerosol relative to formaldehyde and acrolein [2]. The level of acetaldehyde measured within e-cigarette aerosol for our study also reflect those of a prior study identifying that acetaldehyde levels are 10-fold higher than acrolein [34]. Other studies suggest acetaldehyde levels may be at or slightly less than the levels of formaldehyde and acrolein within e-cigarette aerosol [2, 35, 36]. However, acetaldehyde is quite volatile relative to the formaldehyde and acrolein, and the lower levels of acetaldehyde may be attributable to the time for sample preparation causing neutralization and dispersal of acetaldehyde which would account for the lower reading. This is the advantage of using a SIFT MS technique to assess e-cigarette aerosol quantities as the measurements are performed immediately and in real-time. Interestingly, we also observed that the total aldehyde load within e-cigarette aerosol increased with higher percentages of propylene glycol. Together, these findings identify a real-time monitoring system to quantify levels of aldehydes in e-cigarette aerosol identifying that acetaldehyde is a predominant aldehyde within e-cigarette aerosol.

Cigarette smoke-induced oxidative stress within the cardiovascular system is an important trigger for the pathogenesis of cardiovascular diseases, including coronary artery disease, atrial fibrillation, heart failure, and cardiac ischemia-reperfusion injury [15, 37]. E-cigarette use also elevates oxidized low-density lipoprotein, a marker of oxidative stress, 1.5-fold in humans when compared to non-e-cigarette, non-tobacco users [38]. Additionally, e-cigarette aerosol can trigger lipid peroxidation which form endogenous aldehydes including 4-hydroxynonenal (4-HNE) and malondialdehyde (MDA) [27]. As 4-HNE can inhibit ALDH2 enzymatic activity, this can further drive increases in oxidative stress levels from e-cigarette exposures by limiting acetaldehyde metabolism [8]. Importantly, when ALDH2*2 rodents were exposed pair-matched to e-cigarette aerosol exposure with wild type ALDH2 mice, our findings provide new insight that genetics which limit aldehyde metabolism can drive heightened levels of cardiovascular oxidative stress with e-cigarette exposure. This is caused by a lesser capability for the ALDH2*2 enzyme to neutralize reactive species (including reactive oxygen species ROS and endogenous reactive aldehydes) leading to an imbalance requiring less oxidative stress to cause a more pro-reactive environment.

Our study indicates that the ALDH2 enzyme contributes to the heart rate-mediated responses to e-cigarette aerosol. Heart rate increases after e-cigarette or tobacco cigarette use occurs in humans and is correlated with plasma nicotine concentrations [39]. During e-cigarette use, heart rate may increase ~5 beats per minute in humans [40]. We observed that e-cigarette aerosol exposure increases heart rate in ALDH2 and ALDH2*2 rodents, with increases in ALDH2*2 rodents earlier and more pronounced relative to wild type ALDH2 rodents. Taken together, these data indicate that limiting aldehyde metabolism also contributes to the heart rate response of e-cigarettes and likely occurs in a similar mechanism how accumulation of acetaldehyde increases heart rate after alcohol consumption for those carrying the ALDH2*2 variant.

Our study also identifies aldehydes from e-cigarette aerosol can induce oxidative stress. Oxidative stress is known to trigger and exacerbate the pathogenesis of atherosclerosis, hypertension, and diastolic myocardial dysfunction in human volunteers and animal models [41, 42]. Reactive aldehydes from e-cigarette aerosol promote the overproduction of ROS, which exacerbates endogenous lipid peroxidation, protein adduct formation as well as endogenous aldehydes including 4-HNE and MDA, leading to increases in oxidative stress [43]. Kuntic [41] and colleagues identified that reactive aldehydes in the e-cigarette aerosol were important mediators of e-cigarette-induced oxidative stress in wild type mice. Our study confirmed the role of aldehyde metabolism in e-cigarette-induced oxidative stress and importantly identified that inefficient aldehyde metabolism, as for the ALDH2*2 mice, also had higher levels of oxidative stress relative to wild type ALDH2 mice. Other studies [44–46] also reported e-liquid caused the generation of the reactive aldehyde malondialdehyde. In addition, we observed that acetaldehyde as the primary aldehyde in e-cigarette aerosol promotes intracellular Ca^2+^ overload, while the ALDH2*2 variant further enhances the Ca^2+^ influx. Sussan and colleagues showed that elevated ROS production in mice lung induced by the e-cigarette exposure is associated with higher level of lipid peroxidation marker MDA in lung homogenates [27]. We observed similar ROS production and increased levels of lipid peroxidation markers in both wild type ALDH2 mice and ALDH2*2 mice after e-cigarette aerosol exposure. These observations link the aldehyde metabolism to e-cigarette use mediated oxidative stress, calcium handling and eventually cardiovascular dysfunction.

We find that the NF-κB pathway is involved in mediating e-cigarette-induced cardiovascular oxidative stress and at baseline this inflammatory pathway is increased for ALDH2*2 knock-in mice relative to wild type ALDH2 mice. This is secondary to increased baseline levels of ROS in the ALDH2*2 knock-in mice. Similar to prior work this accumulation of intracellular ROS and NF-κB activation by phosphorylation of Ser536 likely leads to downstream activation of pro-inflammatory cytokines which can lead to heighted levels of inflammation [47]. In human pulmonary artery smooth muscle cells, the balance of NF-κB activation and downstream signaling events of inflammation are regulated by ALDH2 [48]. E-cigarette aerosol-induced lung injury has also been linked to oxidative stress and the pro-inflammatory NF-κB pathways [49]. In the context of our present study, ALDH2*2 mice demonstrated a higher level of NF-κB phosphorylation induced from e-cigarette exposure compared with that in wild type ALDH2 mice. Together, this study suggested identifies that activation of NF-κB, which can lead to triggering of a pro-inflammatory pathway is involved in the mechanism how e-cigarettes can elevated levels of cardiovascular oxidative stress.

Although mitochondria are abundant within the heart relative to other organs, ALDH2 expression within the heart is 2.4-fold less relative to the lung and 2.8-fold to the liver. This finding identifies organ systems that are the first exposed to aldehyde sources having a higher capacity to metabolize aldehydes; likely in order to protect the body from oxidative stress caused by inhaled and consumed substances. The tissue specific distribution of ALDH2 puts the cardiovascular system, with less distribution and enzymatic activity relative to other organs, at a greater susceptibility for the reactive aldehyde-induced damage to e-cigarette aerosol which is amplified by carrying the ALDH2*2 genetic variant. Differences in tissue specific expression and activity of ALDH2 may lead to specific organ systems being more susceptible to organ injury from aldehydes [50, 51]. Similar to our findings, a recent non-biased approach to map the proteome of the human body for 32 different organs measured from 14 individuals revealed the liver had ~2-fold higher ALDH2 protein expression relative to the heart [52]. ALDH2 protein expression for our study was also similar between genders (Supplemental Figure 2). Taken together, organ specific levels of ALDH2 expression may be an important player in driving the pathophysiology of e-cigarette aerosol. Understanding ALDH2 expression across organ systems may also advance the understanding and provide a common link to tobacco cigarettes, alcohol, and fried foods, which all are sources of aldehyde exposure, can increase the risk of cardiovascular disease.

Our study should be interpreted within the context of potential limitations. For the rodent portion of the study, we used whole body exposure to e-cigarette aerosol rather than a nose-only exposure. Regardless of the exposure method, our study matched rodents of different ALDH2 genotypes pairwise to e-cigarette exposure to identify what impact the ALDH2*2 variant will have on cardiovascular oxidative stress relative to wild type ALDH2 rodents. Additionally, we used homozygote ALDH2*2 rodents whereas most of the human population are heterozygous for the ALDH2*2 variant. Although we report ALDH2 protein expression was similar between genders, a prior study identified that within the mouse female heart homogenates there was a 20-30% increase in phosphorylation causing a 20-30% increase in enzymatic activity for ALDH2 relative to male heart homogenates [53]. As these gender differences result in a ~10% change in enzymatic activity for wild type ALDH2 rodents this is minor when considering the ALDH2*2 variant triggers a 3-fold lower enzymatic activity relative to the wild type ALDH2 rodents. However, more studies are needed to identify how gender, when coupled with differences in ALDH2 genetics impacts cardiovascular oxidative stress with e-cigarette aerosol exposure. Regardless of these limitations, our data do provide novel insight regarding the potential concerns that within certain populations carrying genetics which limit aldehyde metabolism, oxidative stress is exacerbated with e-cigarette aerosol exposure.

In summary, we identify acetaldehyde as a predominant aldehyde within e-cigarette aerosol and further describe how rodents carrying an inactivating variant of ALDH2, ALDH2*2, are more susceptible to cardiovascular oxidative stress when exposed to e-cigarette aerosol. We also unexpectedly identified that the expression and activity of ALDH2 is lower in the heart relative to the lung and liver which provides a basis to understand how the heart as an organ may be more susceptible aldehyde-induced cardiovascular disease and triggers NF-κB leading to a proinflammatory pathway that leads to elevation of cardiovascular oxidative stress. Ultimately, this study highlights the importance of considering the impact of genetics to develop a further understanding of the health risks of e-cigarette aerosol.

## Supplementary material

Supplementary material is available online

## Author Contribution Statement

X.Y. designed and performed the molecular and cell studies, acquired, analyzed and interpreted the data and wrote the manuscript. X.Z. and P.S. designed the animal e-cigarette exposure. X.Z. and F.X. conducted the animal e-cigarette exposure studies, and X.Z wrote the method for mice e-cigarette exposure. R.C. conducted the e-cigarette aerosol collection and quantification studies, and wrote the method for the aerosol analysis. F.X. conducted the cardiomyocytes isolation. E.R.G. design the study, interpreted the data, reviewed and edited manuscript, and supervised the study.

