## supplemental tables & figures for "E-cigarette aerosol exacerbates cardiovascular oxidative stress in mice with an inactive aldehyde dehydrogenase 2 enzyme"

Short Title: E-cigarette aerosol, ALDH2*2 and oxidative stress

**Address Correspondence to**:

Eric R. Gross, MD, PhD,
Department of Anesthesiology, Perioperative and Pain Medicine
School of Medicine,
Stanford University, Stanford, CA. 94305


### SUPPLEMENTAL MATERIAL

**Supplemental Table 1.**

| Metabolite | Reagent Ion | Reaction Ratio | Mass (m/z) |
|---|---|---|---|
| Acetaldehyde | $H_3O^+$ | $3.70 \times 10^{-9}$ | 45, 63 |
| | $NO^+$ | $6.00 \times 10^{-9}$ | 43, 61 |
| Formaldehyde | $H_3O^+$ | $3.40 \times 10^{-9}$ | 31, 49 |
| Acrolein | $H_3O^+$ | $4.20 \times 10^{-9}$ | 57, 75 |
| | $NO^+$ | $1.40 \times 10^{-9}$ | 55, 86 |
| Nicotine | $H_3O^+$ | $3.0 \times 10^{-9}$ | 163 |
| | $NO^+$ | $2.5 \times 10^{-9}$ | 162 |
| | $O_2^+$ | $1.8 \times 10^{-9}$ | 162 |

**Supplemental Table 1.** Reagent ions, reaction ratio and mass used for SIFT-MS to detect and identify nicotine and aldehydes within e-cigarette aerosol.

**Supplemental Figure 1**

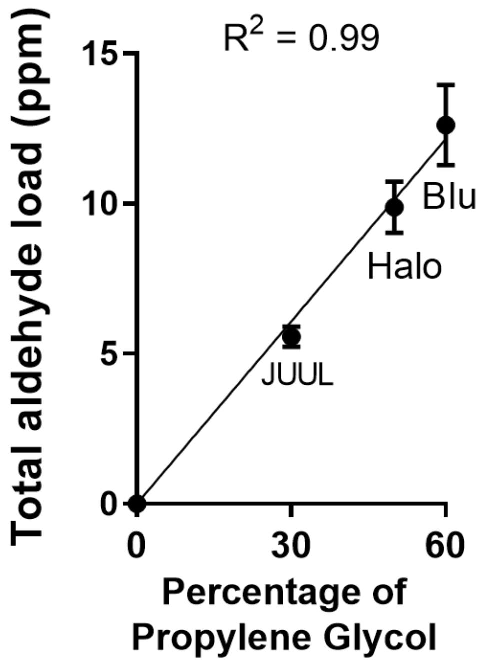


**Supplemental Figure 1**. Total aldehyde load (ppm) relative to the percentage of propylene glycol from different e-cigarette brands. $R^2$ was calculated by Pearson correlation. n= 20 replicates per e-cigarette.

**Supplemental Figure 2**

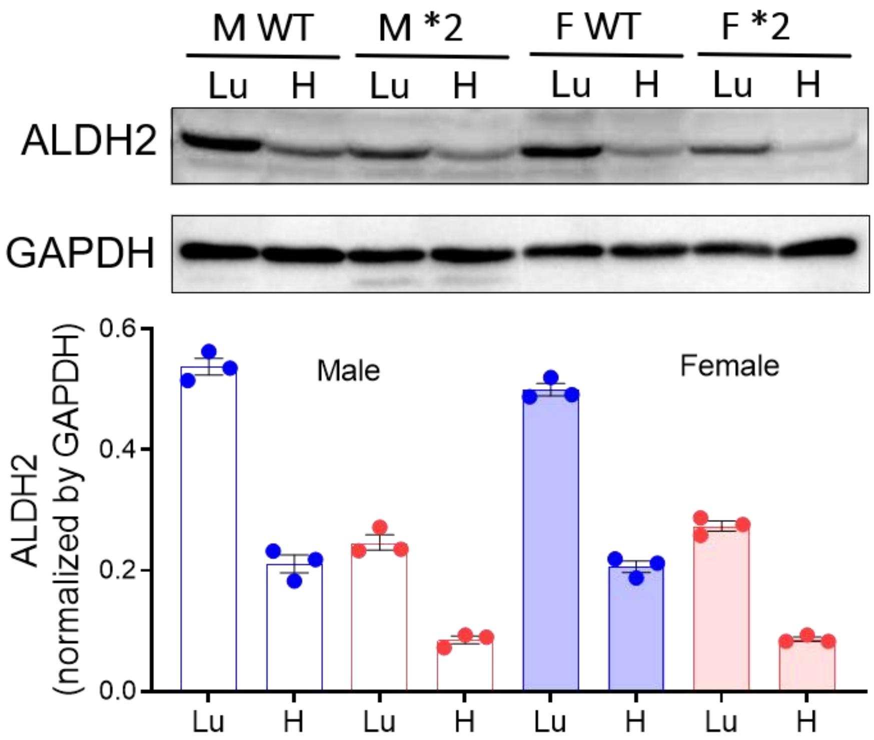


**Supplemental Figure 2. Western blot of male and female wild type ALDH2 and ALDH2*2 rodents.** ALDH2 protein expression is gender independent for both wild type ALDH2 and ALDH2*2 rodents.

**Supplemental Figure 3**

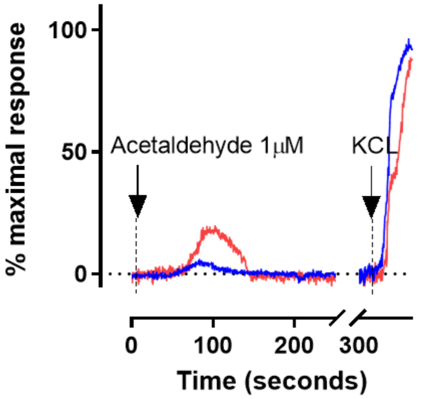


**Supplemental Figure 3. Calcium response in primary cardiac myocytes exposed to acetaldehyde.** Calcium influx induced by 1 µM acetaldehyde (n = 21 cells for ALDH2 and n= 20 cells for ALDH2*2 from 4 biological replicates) *p<0.05 ALDH2*2 versus wild type ALDH2.

**Supplemental Figure 4**

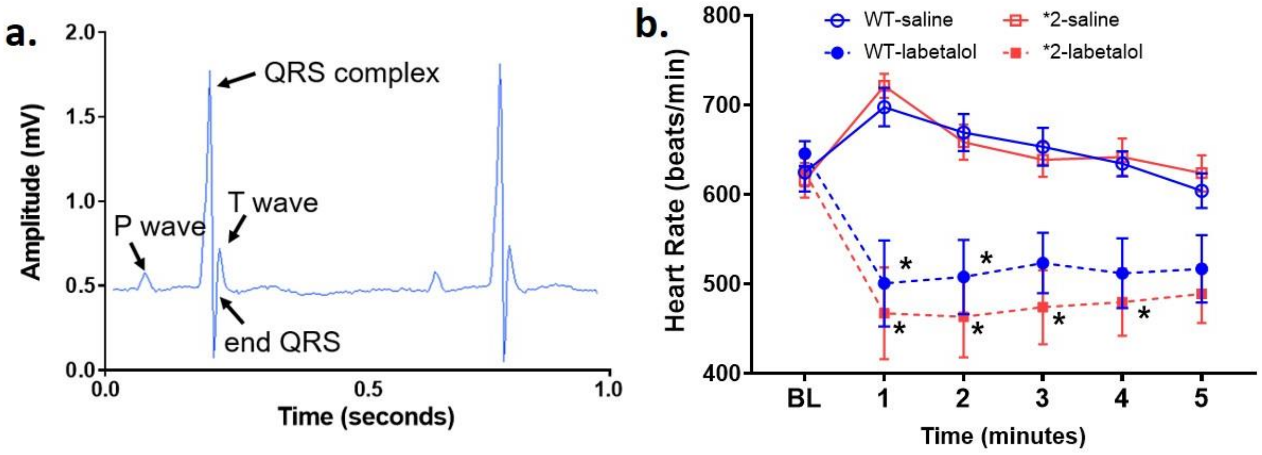


**Supplemental Figure 4**. **EKG telemeter testing with labetolol.** ***a)***. Representative EKG waveform captured from mouse telemeter. Arrows show P wave, QRS complex and T wave respectively. ***b)***. Time course for mean heart rate before and after intraperitoneal injection of labetolol or saline. Data was expressed as mean ± SEM. *p<0.05 saline injection versus labetolol injection.

**Supplemental Figure 5**

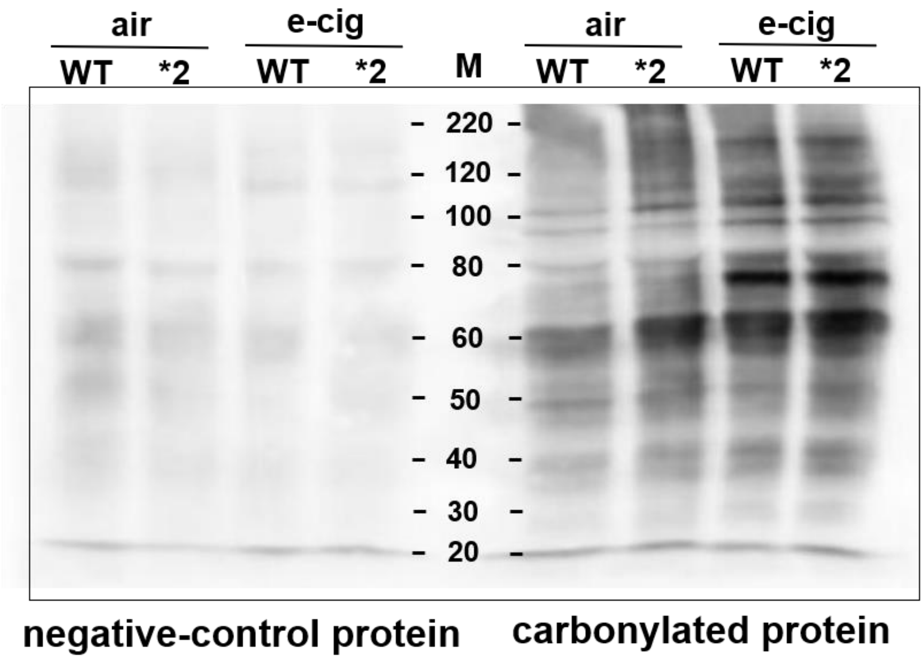


**Supplemental figure 5. Protein carbonylation blot.** Uncut blot of the negative control (left half of panel) and DNP-derivatized protein carbonyls (right half of panel).
